# Sensory-thresholded switch of neural firing states in a computational model of the ventromedial hypothalamus

**DOI:** 10.1101/2021.12.02.470929

**Authors:** Ryan Rahy, Hiroki Asari, Cornelius T. Gross

**Affiliations:** Epigenetics & Neurobiology Unit, European Molecular Biology Laboratory (EMBL), Via Ramarini 32, 00015 Monterotondo (RM), Italy

**Author notes:** Corresponding author: Cornelius T. Gross.

## Abstract

The mouse ventromedial hypothalamus (VMH) is both necessary and sufficient for defensive responses to predator and social threats. Defensive behaviors typically involve cautious approach toward potentially threatening stimuli aimed at obtaining information about the risk involved, followed by sudden avoidance and flight behavior to escape harm. *In vivo* neural recording studies in mice have identified two major populations of VMH neurons that either increase their firing activity as the animal approaches the threat (called *Assessment*+ cells) or increase their activity as the animal flees the threat (called *Flight*+ cells). Interestingly, *Assessment*+ and *Flight*+ cells abruptly decrease and increase their firing activity, respectively, at the decision point for flight, creating an escape-related “switch” in functional state. This suggests that the activity of the two cell types in VMH is coordinated and could result from local circuit interactions. Here, we used computational modelling to test if a local inhibitory feedback circuit could give rise to key features of the neural activity seen in VMH during the approach-to-flight transition. Starting from a simple dual-population inhibitory feedback circuit receiving repeated trains of monotonically increasing sensory input to mimic approach to threat, we tested the requirement for balanced sensory input, balanced feedback, short-term synaptic plasticity, rebound excitation, and inhibitory feedback exclusivity to reproduce an abrupt, sensory-thresholded reciprocal firing change that resembles *Assessment*+ and *Flight*+ cell activity seen *in vivo*. Our work demonstrates that a relatively simple local circuit architecture is sufficient for the emergence of firing patterns similar to those seen *in vivo* and suggests that a reiterative process of experimental and computational work may be a fruitful avenue for better understanding the functional organization of mammalian instinctive behaviors at the circuit level.

## Introduction

A major goal of current neuroscience research is to arrive at a circuit-based understanding of behavior so as to be able to better describe and categorize the origins of animal behavior and rationally design behavior-modifying treatments. The last decade has seen an enormous advance in the tools available to behavioral circuit neuroscientists who can now record hundreds of neurons simultaneously in freely behaving laboratory animals and activate or inhibit the firing of defined classes of neurons at the millisecond timescale. [1] These tools have ushered in an era in which new approaches are needed to understand the link between neural firing activity and behavior, and there has been a renewed call to engage with theoretical neuroscientists to build predictive models and derive testable hypotheses. Ideally, such collaborations between experimentalists and theoreticians would fuel a virtuous experiment-theory cycle that could help arrive at an understanding of the circuit basis of behavior. [2,3]

The medial hypothalamic defensive system offers a fertile subject for circuit behavior neuroscience because it consists of a series of interconnected nuclei that are anatomically and functionally conserved across mammals, including primates, [4,5] and it forms a relatively simple subcortical pathway linking sensory input to motor output. [4–6] Extensive work has been carried out on a central node in this system, the ventromedial hypothalamus (VMH), and inhibition of this nucleus blocks defensive responses to predator and social threats, [7,8] while activation elicits flight behavior. [8–13] Consistent with its anatomical conservation between rodents and primates [5,12] electrical stimulation of VMH in humans elicits intense arousal, negative emotion, and panic, [14] supporting the hypothesis that VMH encodes an internal defensive state that is relevant to understanding instinctive fear in humans. [15]

*In vivo* single unit electrophysiology and calcium endoscopy recordings in freely behaving mice have revealed that VMH encodes both sensory and motor features of defensive behavior. [10,16] Single unit electrophysiology recordings were carried out in VMH as mice approached and then fled a live rat, a natural predator. Twenty-eight percent of units showed significant changes in firing during the approach-to-flight behavior. Of these, 39% showed either increased or decreased firing during approach – called *Assessment*+ and *Assessment*-cells, respectively – while 61% showed either increased or decreased firing during flight – called *Flight*+ and *Flight*-cells, respectively. [16] Notably, *Assessment*+ cells abruptly decreased firing at the moment in which the animal initiated escape, while *Flight*+ cells abruptly increased firing at this point. These data demonstrate that separate neuron populations in VMH encode sensory information about threat and motor information about the defensive behavioral outcome and suggest that a transition from sensory to motor encoding occurs immediately prior to flight. Moreover, because the firing rate of *Assessment*+ cells increased linearly as the animal approached the threat [16] the abrupt change in firing of *Assessment*+ and *Flight*+ cells appeared to occur in a manner whose probability was increased by or thresholded to the intensity of threat exposure (**Fig S1**). While these recordings were carried out in the dorsomedial VMH (VMHdm) during exposure of mice to a rat, similar *Assessment*+ and *Flight*+ cells were seen in the neighboring ventrolateral VMH (VMHvl) during exposure of mice to a social aggressor, [10] consistent with the anatomical segregation of predator and social threat processing [6,7,17,18] and suggesting a generalized sensory-motor encoding of innate defensive responses across threats in the medial hypothalamus.

Two previous studies have reported computational models of VMH. [19,20] The former was based on a classification of neurons with putative differences in intrinsic excitability based on hazard plots of firing propensity extracted from *in vivo* single unit recordings of VMHdm neurons in anesthetized rats. [21] These excitability properties were modelled in a network of interconnected excitatory neurons to derive network firing activities. The study concluded that slow oscillatory rhythms and network bistability similar to those experimentally observed could emerge from a model of a heterogeneous population of VMH neurons. Interestingly, the model revealed subclasses of cells that showed spontaneous ON/OFF firing states, pointing to an intrinsic propensity for sudden state transitions among VMH neurons. [19] The second study constructed a circuit model of the mouse VMHvl composed of interconnected excitatory core neurons receiving sensory input and inhibitory feedback. [20] The study aimed to identify conditions under which persistent neural firing activity emerged in response to transient sensory input. Such persistent activity was proposed as a substrate for an internal defensive state in VMH deriving from experimental [8] and theoretical [22] considerations. The model showed that the addition of slow-acting neuromodulatory transmission between VMH core neurons led to sustained bulk activity of core neurons. Interestingly, in addition to sensory-ON neurons whose activity matched sensory input, neurons with sensory-OFF properties were observed that showed peak activity at stimulus offset, demonstrating that the combination of excitatory interconnectivity and feedback inhibition could give rise to neuron classes with reciprocal sensory-ON/OFF stimulus responses.

We explored whether a computational model of local VMH circuitry could give rise to the transition in firing state observed during approach-to-flight behavior [10,16] and be used to constrain future experimental circuit manipulation studies. We aimed to produce a parsimonious model in which three firing features would emerge: 1) distinct *Assessment+* and *Flight+*-like cell populations, 2) a sensory-input thresholded abrupt change in global firing state, and 3) a simultaneous and opposite change in firing activity of *Assessment+* and *Flight+*-like cells at threshold. In other words, a reciprocal ‘switch’ in activity of Assessment and Flight cell populations should occur when sensory input passes a specific level. Our findings show that thresholded switching between Assessment and Flight-like populations can occur under conditions of excitatory neuron interconnectivity coupled to feedback inhibition with neuromodulatory rebound excitation, a feature observed in *in vivo* neural recordings. [23] These conditions set out several non-obvious hypotheses for experimental testing using circuit recording and manipulation tools in behaving animals.

## Methods

### Circuit model

To identify key network features underlying a sensory-thresholded switch between *Assessment*+ and *Flight*+ cells we built circuit models in NEST [24] consisting of three major cell types (Amygdala, VMH core, VMH shell) and introduced different network geometries and properties among them (Models 1-4; model simulation scripts are available at https://gitlab.com/gross-group/vmh-models). All models used leaky integrate-and-fire neurons with exponentially-shaped postsynaptic currents [25] and Bernoulli synapses (static synapses with release probability = *p*_*release*_), with a timestep of 0.1 ms. All connections within and between populations followed a Bernoulli distribution (connection probability = *p*_*connection*_). VMH core included two populations of excitatory neurons (N = 100 each, Assessment and Flight neurons). Parameters were chosen based on previously reported *in vitro* slice electrophysiological properties of VMH core and shell neurons (core: resting membrane potential = −58.6 mV, action potential threshold = −43.1 mV, membrane time constant = 40.3 ms; shell: resting membrane potential = −56.5 mV, action potential threshold = −42 mV, membrane time constant = 41.6 ms). [26] VMH shell included either two populations (Models 1-3: N = 20 each, exclusive Assessment-to-Flight/Flight-to-Assessment connectivity) or a single population (Model 4: N = 40 neurons, random Assessment-to-Flight/Flight-to-Assessment connectivity). Shell neurons are predisposed to having higher firing rates [26] and were given higher noise frequencies than excitatory core neurons (core: 5 Hz; shell: 10 Hz). Where relevant, imbalanced weights of shell-to-Assessment and shell-to-Flight connections were set by multiplying the inhibitory weight of selected connections by a ratio 1 ≤ *r*_*feedback*_ ≤ 20 (Model 1). The Amygdala consisted of a uniform population of non-interconnected excitatory neurons (N = 100) receiving sensory input and asymmetrically projecting onto both VMH core populations with varying connection density to Flight neurons obtained by multiplying the Amygdala-to-Flight *p*_*connection*_ by a ratio 0 ≤ *r*_*input*_ ≤ 1 (Models 1-4). Weights of Amygdala connections to VMH core were sampled randomly from a normal distribution, while delay values were sampled from a uniform distribution to introduce variability in the input received by VMH core. All parameter values used are listed in **Table S1**. We aimed to make these values as consistent as possible across models, but due to differing model features and synaptic properties some variation was necessary.

### Sensory input

To simulate the sensory input that a mouse experiences during approach-to-flight we employed an inhomogeneous Poisson process that generated spiking signals to the Amygdala population based on a time-varying firing rate function *λ*(*t*). In each trial (20 s) we used *λ*(*t*) = max(*f*_*baseline*_, *a*_*model*_ *t*) for the first half of the trial (0 < *t* < 10 *s*) to approximate the increasing sensory signal during approach behavior (assessment phase), and *λ*(*t*) = max(*f*_*baseline*_, *f*_*max,model*_ − 4*a*_*model*_(*t* − 10)) for the remaining 10 s (10 < *t* < 20 *s*) to simulate the decreasing sensory signal during avoidance behavior (flight phase), where *f*_*baseline*_ = 40 *Hz* sets the baseline level of input to the Amygdalar cells, *f*_*max,model*_ represents the maximum sensory input firing rate for each model, and 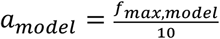 is the slope for each model. The values for each *f*_*max,model*_ can be found in **Table S1**. Note that a mouse typically flees faster than it approaches a threat, hence the rapid drop in sensory input (**Fig S1**). [16] We included multiple trials in a single simulation run to assess consistency in the network response without resetting parameters. Each run of our simulation included 10 consecutive trials with inter-trial intervals of 6 s and buffer periods of 2 s at the beginning and end. During the inter-trial intervals and buffer periods we used *λ*(*t*) = *f*_*baseline*_ to represent a baseline level of sensory input to the Amygdala.

### Synaptic plasticity and rebound

Short-term plasticity was implemented in core-efferent synapses in Model 2 as described in the Tsodyks-Markram synaptic model. [25] Briefly, this model reproduces the effect of plasticity by increasing the release probability of a given synapse by a certain increment with each presynaptic spike (facilitation time constant, 0 ≤ τ_*facilitation*_ ≤ 10000 *ms*, reflects how fast the synapse returns to the initial release probability). It also decreases the synaptic resources after each spike (depression time constant, 0 ≤ τ_*depression*_ ≤ 10000 *ms*, reflects the rate of synaptic vesicle replenishment). Because NEST does not allow for having both fast and slow transmission in the same synapse, excitatory rebound was introduced into Models 3 and 4 by adding an intermediate population of neurons between shell and core with non-probabilistic release synapses and a slow excitatory effect (delay time, 0 ≤ *t*_*delay*_ ≤ 50 *ms*; rebound time constant, 0 ≤ τ_*rebound*_ ≤ 50 *ms*; estimated from experimental data: *t*_*delay*_ = 14 *ms*, τ_*rebound*_ = 10 *ms*). [23]

### Model feature analysis

Parameter exploration of model-specific features was performed by running each model over single-trial-per-run simulations with different values for the key parameters of each model within specified ranges. Each simulation was then classified qualitatively; a simulation was classified as a ‘Switch’ if Assessment and Flight firing patterns satisfied the following criteria: 1) Assessment onset was gradual, decrease was sudden, 2) Flight onset was sudden, decrease was gradual, 3) Firing was reciprocal – Flight onset reduced Assessment firing to at least 40% of maximum, 4) Signals were sustained, but Flight firing returned to baseline within trial time, and 5) Assessment firing did not go up beyond baseline levels after Flight firing stopped. Simulations that did not satisfy all 5 criteria were classified as an ‘Intermittent reciprocal firing simulation’ where Assessment and Flight neurons showed unstable, repeated reciprocal changes in firing activity, or ‘None’ when the simulation could not be placed in any other category. ‘None’ simulations were not represented in the scatterplots of the parameter space. To test for thresholding of sensory input, we ran each model over simulations with 10 trials per run and varying slopes of sensory input. For each trial, the latency was calculated as the time from the start of the increase in sensory input to the onset of the switch. An inverse correlation between latency and sensory input at the onset of the switch was interpreted as a sign of switch thresholding to a certain level of sensory input.

### Analysis of firing activity

All analyses of neuron firing were carried out in Python using the PySpike package. [27] For population analysis we first computed the peri-stimulus time histogram (PSTH; bin size = 20 ms) of all cells combined for each population (e.g. all Assessment or all Flight neurons). For each of the 10 trials we determined the onset time of the switch by identifying the moment at which the firing of the Flight population surpassed that of the Assessment population by more than 25% of the overall PSTH maximum of both populations. Finally, we averaged maximum-normalized 20-s windows of each population PSTH centered around each putative switch. For single neuron analysis we repeated the same procedure using the firing activity of individual cells.

## Results

### Circuit Model Architecture

We argued that a successful computational model of VMH should lead to the emergence of two distinct neuron populations showing an abrupt reciprocal change in activity at a sensory input threshold – corresponding to *Assessment+* and *Flight+* neuron activity reported during approach-to-flight behavior (**Fig S1**). [10,16] To explore the connectivity, intrinsic excitability, and synaptic plasticity conditions under which such a sensory-thresholded, reciprocal neuronal response might emerge, we implemented a circuit model of leaky integrate-and-fire neurons in the NEST computing environment. [24] The VMH consists of a mainly excitatory core with a predominantly inhibitory shell, with synaptic connections between and within core and shell neurons. [20,26,28] Our model therefore consisted of a group of interconnected excitatory core VMH neurons (N = 200) segregated into two populations, *a priori* called Assessment and Flight cells (N = 100 each), so as to be able to vary their physiological characteristics separately, if needed. These neurons received reciprocal inhibitory feedback via shell VMH inhibitory neurons (N = 40) and sensory input from excitatory Amygdala neurons (N = 100). Previous work showed that amygdala inputs to VMH convey predator and social threat information, [29,30] including pheromone/kairomone-based information about threat identity [11,31–33] and polymodal information about threat intensity. [34–38] Because *Assessment+*, but not *Flight*+ neuron firing *in vivo* correlated linearly with inverse distance to threat – and thus to the intensity of threat-related sensory input [16] – we explored both the simple condition under which Assessment, but not Flight cells received sensory input from Amygdala, and also conditions under which Assessment and Flight cells received varying relative Amygdala inputs (Flight = 0 to 100% Assessment input; see **Methods**). Throughout our study sensory input aimed at representing approach-to-flight behavior was modeled by repeated trains of brief, linearly increasing, and noisy excitatory input to Amygdala cells (**Fig S2**).

### Model 1: Asymmetric feedback inhibition

First, we explored how the balance of feedback inhibition onto Assessment and Flight cells affected VMH core neuron responses to sensory input (Model 1, **Fig 1a**). A systematic assessment of core neuron sensory responses under conditions of varying Amygdala input density (0 ≤ *r*_*input*_ ≤ 1) and feedback inhibition strength (1 ≤ *r*_*feedback*_ ≤ 20) revealed parameters under which a simultaneous and opposite change in firing activity of Assessment and Flight neuron sensory responses emerged – a condition we refer to as a ‘switch’ in activity – when Assessment cells received denser sensory input (*r*_*input*_ ∈ [0.2, 0.8]) and stronger feedback inhibition (*r*_*feedback*_ ∈ [2, 20]) than Flight cells. This finding argues that intrinsic differences in the connectivity of VMH cell populations may be necessary for differential Flight and Assessment responses to sensory input (**Fig 1b, S3a**; representative simulation: *r*_*input*_ = 0.5, *r*_*feedback*_ = 2.5, **Fig 1cd**). Importantly, the switch in Assessment and Flight neuron firing occurred during the increasing phase of sensory input suggesting that the switch may be thresholded to a specific level of sensory input. Consistent with such sensory input thresholding, the latency to switch onset varied inversely with sensory input slope (**Fig 1e**).

**Figure 1.**
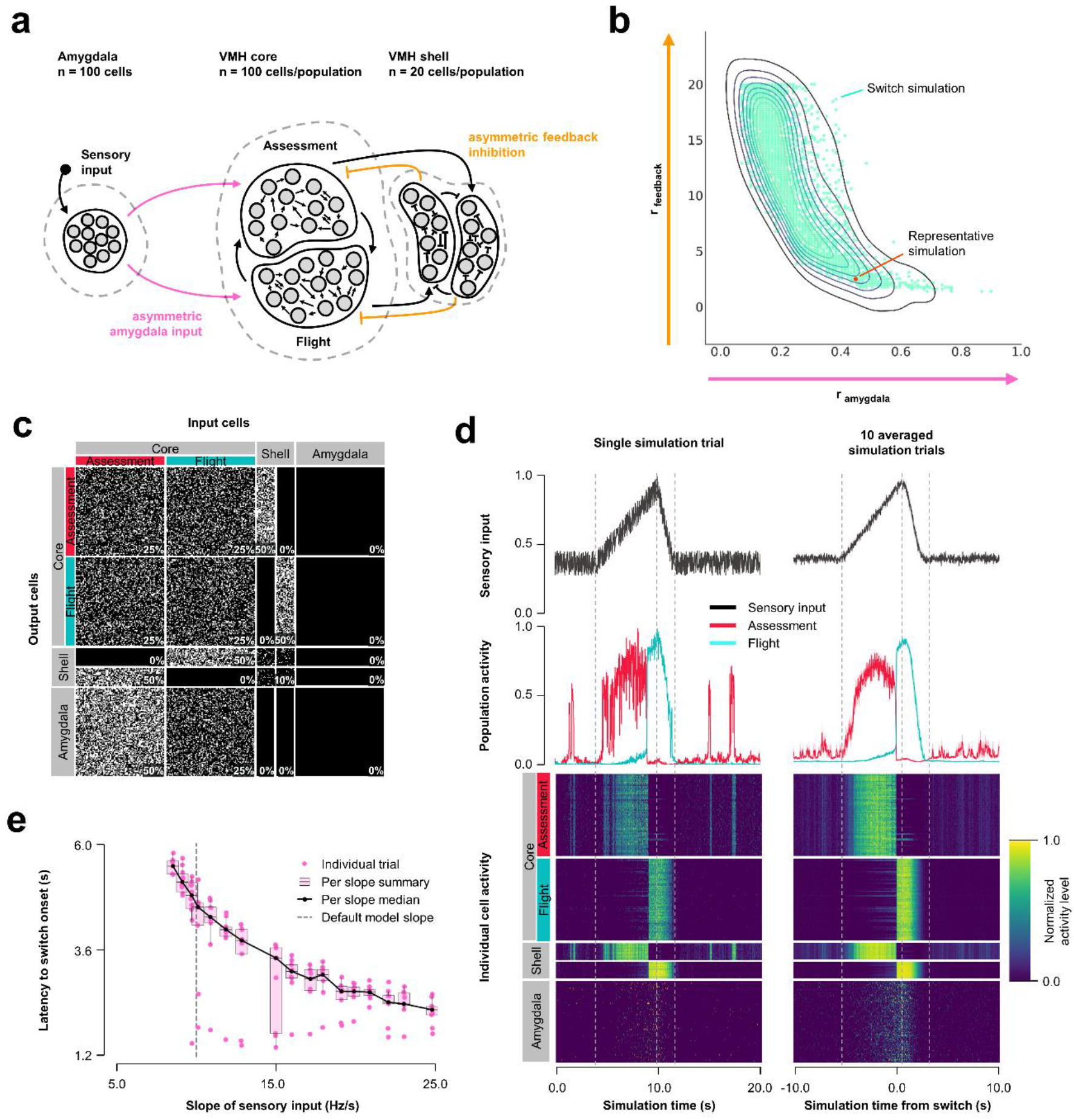
Model 1: Asymmetric feedback inhibition. (**a**) Circuit structure in which both the ratio (*r*_*input*_) of the probability of Amygdala-Flight/Amygdala-Assessment connectivity (pink) as well as the ratio (*r*_*feedback*_) of the weights of shell-Assessment/shell-Flight connectivity (orange) were systematically varied. (**b**) Systematic model exploration identified parameters that satisfy criteria for Assessment/Flight switching (turquoise, see **Methods**), where only simulation parameters that satisfy these criteria are shown. A representative switch simulation is highlighted in orange. (**c**) Network connectivity matrix of the representative simulation showing connectivity probabilities between all neuron populations. (**d**) Network firing activity from the representative simulation for a single-trial simulation run (left) or a 10-trial averaged simulation run (right). Top row shows sensory input, middle row shows firing patterns of Assessment (red) and Flight (blue) populations, and bottom row shows firing activity of all individual cells, separated by population. All trials were centered around the time the switch occurs before averaging. The vertical dashed lines mark the beginning and end of the assessment and flight phases of the sensory input (see **Methods**). (**e**) Plot of sensory input slope vs. latency of switching showing an inverse relationship between latency and sensory threshold. Only slope values for which a switch was observed are indicated.

### Model 2: Short-term plasticity

Because Assessment/Flight responses in Model 1 remained relatively gradual compared to the abrupt reciprocal change in firing rates seen *in vivo* (**Fig S1**) we explored whether the inclusion of synaptic plasticity might convey a sharpening of the switch. Short-term synaptic plasticity can have a presynaptic or postsynaptic origin and is a short-lasting non-linear neurotransmission mechanism described across many neuronal connections [39] including amygdala inputs to VMH. [37] We added short-term, pre-synaptic, facilitating plasticity to all core neuron outputs, including both core-core and core-shell synapses, in a model with varying Amygdala input density and symmetric feedback inhibition strength and examined Assessment and Flight neuron responses (Model 2, **Fig 2a**). A systematic exploration of synaptic plasticity parameters (0 ≤ τ_*facilitation*_ ≤ 10000 *ms*; 0 ≤ τ_*depression*_ ≤ 10000 *ms*) and Amygdala input asymmetry (0 ≤ *r*_*input*_ ≤ 1) failed to identify parameters that gave rise to a stable switch in Assessment and Flight neuron responses (**Fig 2b**; representative simulation: τ_*facilitation*_ = 3158 *ms*, τ_*depression*_ = 527 *ms, r*_*input*_ = 0.5, **Fig 2cd**). Under these conditions Assessment neurons showed sensory-ON responses but Flight neurons exhibited intermittent firing responses that failed to show a consistent pattern across trials. Nevertheless, the introduction of short-term synaptic plasticity was able to sharpen reciprocal Assessment/Flight neuron activity changes as Flight neuron firing was associated with rapid suppression of Assessment neurons. Interestingly, unlike in Model 1, this behavior was independent of Amygdala input asymmetry (**Fig S3b**) showing that abrupt reciprocal Assessment/Flight firing changes can occur even when they have indistinguishable sensory inputs.

**Figure 2.**
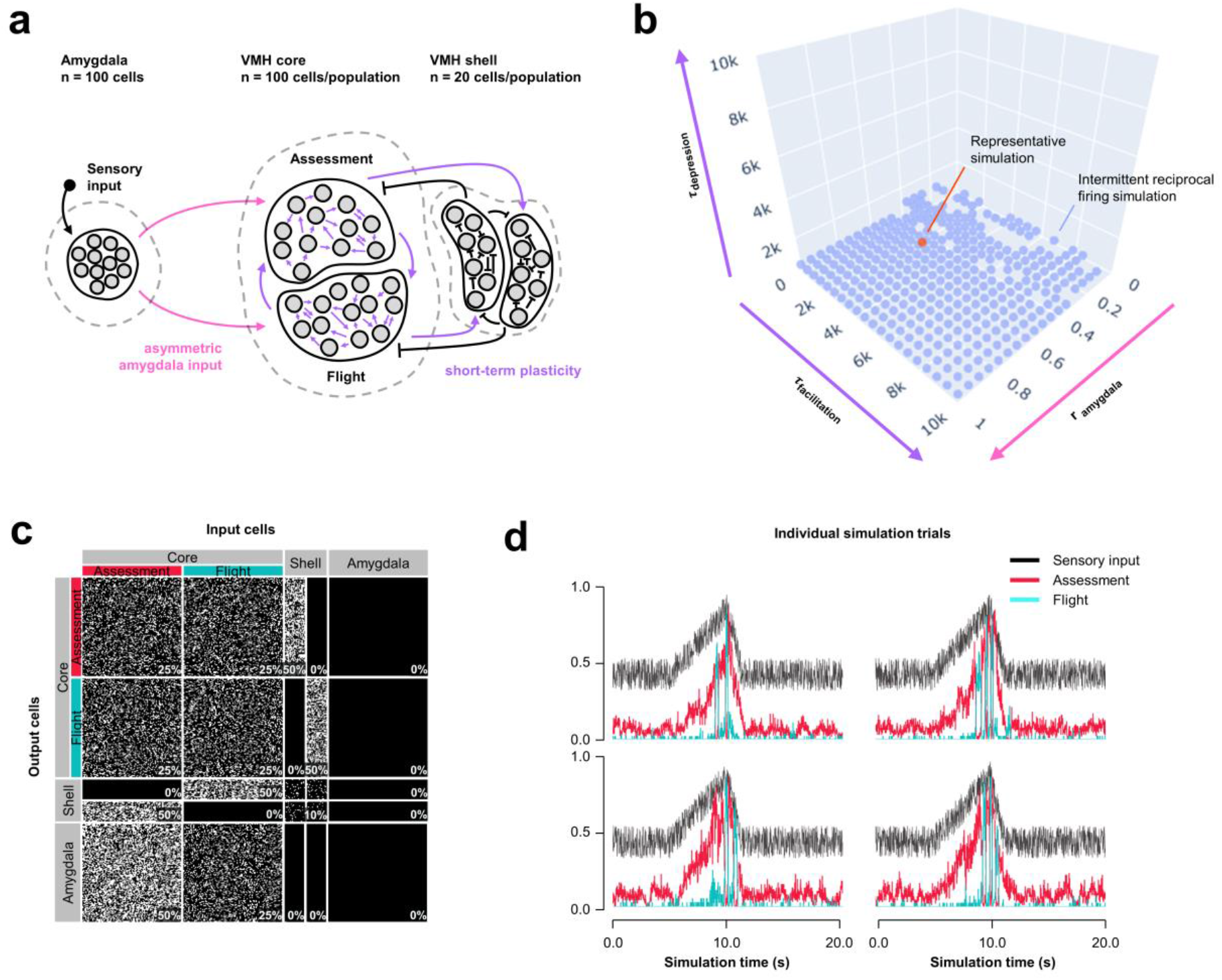
Model 2: Short-term plasticity. (**a**) Circuit structure in which both depression and facilitation time constants for short-term plasticity introduced at all excitatory core output connections (purple) as well as the ratio (*r*_*input*_) of the probability of Amygdala-Flight/Amygdala-Assessment connectivity (pink) were systematically varied. (**b**) Systematic model exploration failed to identify parameters that consistently produced a stable switch. Instead, model parameters that showed intermittent reciprocal Assessment/Flight activity are indicated (blue), as is a representative reciprocal simulation (orange). (**c**) Network connectivity matrix of the representative simulation showing connectivity probabilities between all neuron populations. (**d**) Network firing activity from the representative simulation of a single 10-trial simulation run showing six individual trials with unstable and inconsistent firing patterns.

### Model 3: Excitatory rebound

Next, we explored whether other non-linear synaptic mechanisms might facilitate more stable, but nevertheless abrupt, reciprocal activity changes in response to sensory input. VMH neurons have been shown to exhibit slow-acting excitatory rebound responses to the optogenetic activation of inhibitory inputs. [23] Importantly, such rebound responses were restricted to neurons that showed decreased (called *Attack*-neurons), but not increased (called *Attack*+ neurons) firing during approach-to-attack behavior. [23] Although in this study the firing of *Attack*+/-neurons was not investigated during approach-to-flight behavior, we speculate that these *Attack*+ and *Attack*-cells correspond to *Assessment*+ and *Flight*+ cells, respectively, because of their similar firing activity responses during social approach (*Flight*+ cells frequently showed decreased firing during approach). [10,16] Based on these considerations we incorporated slow-acting excitatory rebound responses exclusively at inhibitory inputs onto Flight cells (Model 3, **Fig 3a, S4a**). A systematic exploration of excitatory rebound parameters (0 ≤ *t*_*delay*_ ≤ 50 *ms*; 0 ≤ τ_*rebound*_ ≤ 50 *ms*) and Amygdala input density asymmetry (0 ≤ *r*_*input*_ ≤ 1) identified a relatively narrow range at which Assessment and Flight neurons showed reliable and stable switching behavior (**Fig 3b**; representative simulation: *t*_*delay*_ = 14 *ms*, τ_*rebound*_ = 10 *ms, r*_*input*_ = 0.7; **Fig 3cd, S4c**). Stable switching was seen regardless of delay time, but only within a narrow range of rebound time constant (τ_*rebound*_ ∈ [7.9,23.7] *ms*) that matched well to that observed experimentally. [23] Amygdala input to Flight cells contributed to stabilizing the response dynamics and avoiding intermittent firing, reinforcing the observation that symmetric Amygdala inputs to VMH core neurons is compatible with the emergence of reciprocal firing responses (**Fig S3c**). Again, Model 3 exhibited sensory input thresholding with latency to switch onset varying inversely with sensory input slope (**Fig 3e**).

**Figure 3.**
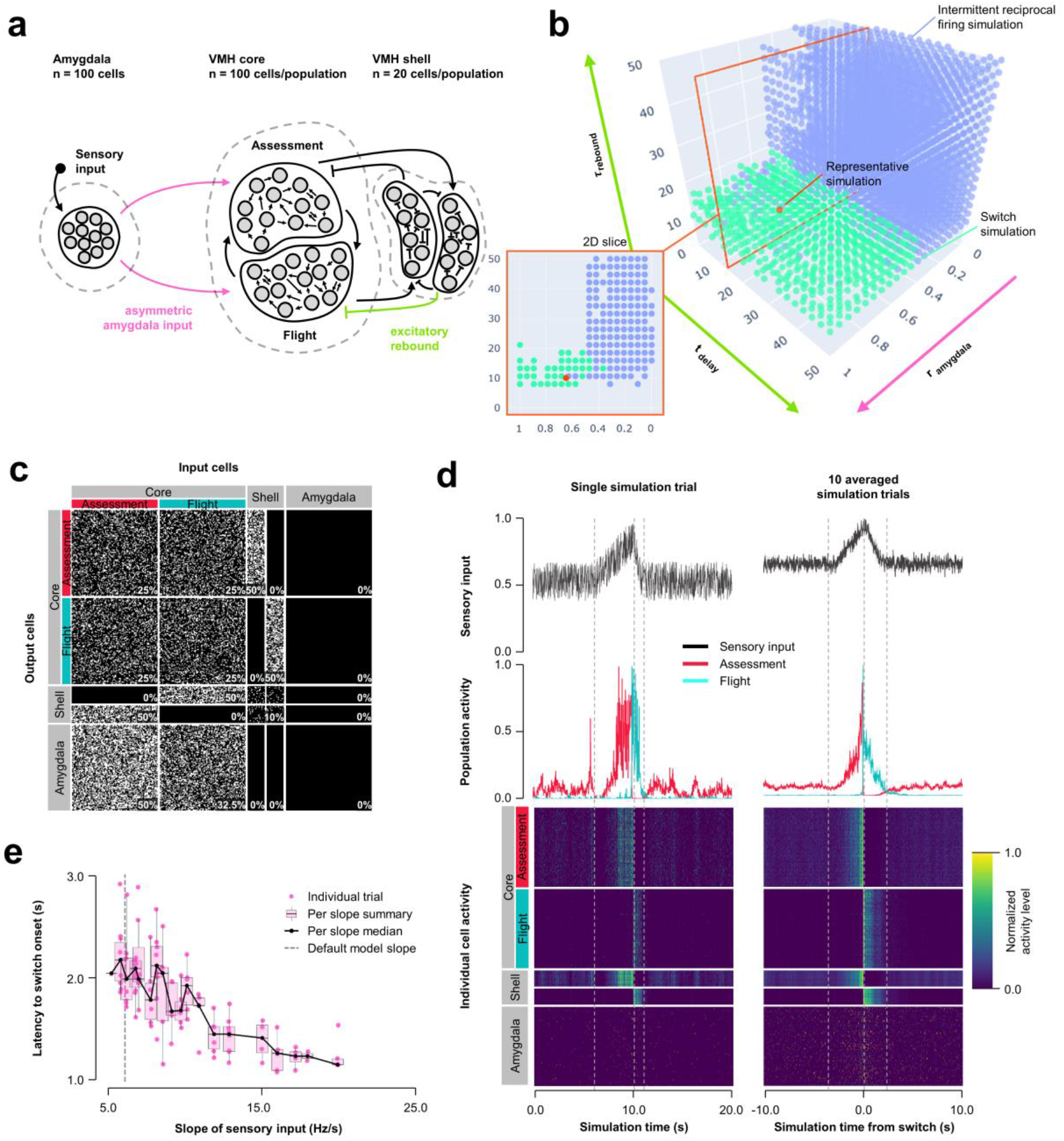
Model 3: Excitatory rebound. (**a**) Circuit structure in which both delay and membrane time constants for excitatory rebound in inhibitory inputs onto Flight neurons (green) as well as the ratio (*r*_*input*_) of the probability of Amygdala-Flight/Amygdala-Assessment connectivity (pink) were systematically varied. (**b**) Systematic model exploration identified parameters that showed consistent Assessment/Flight switching (turquoise, see **Methods**) or intermittent reciprocal firing (blue). Parameter combinations not falling into either category are not shown. A selected representative switch simulation is indicated in orange, and the plane in the parameter space containing this simulation (at *t*_*delay*_ = 14 *ms*, also in orange) is shown in the 2D slice in the inset on the bottom left. (**c**) Network connectivity matrix of the representative simulation showing connectivity probabilities between all neuron populations. (**d**) Network firing activity for the representative simulation from a single-trial simulation run (left) or a 10-trial averaged simulation run (right). Top row shows sensory input, middle row shows firing patterns of Assessment (red) and Flight (blue) populations, and bottom row shows firing activity of all individual cells, separated by population. All trials are centered around the time the switch occurred before being averaged. The vertical dashed lines mark the beginning and end of the assessment and flight phases of the sensory input (see **Methods**). (**e**) Plot of sensory input slope vs. latency of switching showing an inverse relationship between latency and sensory threshold. Note: only slope values for which a switch was observed are indicated.

### Model 4: Non-exclusive feedback inhibition

Finally, we examined an alternative connectivity for feedback inhibition. In Models 1-3, feedback inhibition was exclusive, with each shell neuron receiving input from either Assessment or Flight neurons and subsequently providing feedback to Flight or Assessment neurons, respectively (**Fig 1–3a**). Such exclusive feedback wiring would require high fidelity labelled-line mapping for core-shell-core feedback and could be considered unlikely to occur in nature. In Model 4 we explored an alternative architecture in which core and shell neurons were randomly connected and shell neurons formed a single population (**Fig 4a, S4b**). Systematic exploration of parameter space in this non-exclusive connectivity model using the same conditions as in Model 3 revealed stable Assessment/Flight neuron switching in a limited rebound parameter space (**Fig 4b**; τ_*rebound*_ ∈ [1.9,15.9] *ms, t*_*delay*_ ∈ [4.5,16.5] *ms*; representative simulation: *t*_*delay*_ = 14 *ms*, τ_*rebound*_ = 10 *ms, r*_*input*_ = 0.7; **Fig 4cd, S4d**). Critically, Assessment/Flight neuron switching showed properties similar to those seen *in vivo*, including thresholding and abrupt onset. Moreover, switching persisted even when Amygdala inputs to Assessment and Flight neurons were weighted equally (*r*_*input*_ = 1; **Fig S3d**) and showed sensory input thresholding (**Fig 4e**). These findings demonstrate that the expression of slow-acting excitatory rebound exclusively in Flight neurons is sufficient for the emergence of distinct Assessment and Flight neurons with thresholded, reciprocal sensory responses in the absence of any differences in their connectivity or intrinsic response properties.

**Figure 4.**
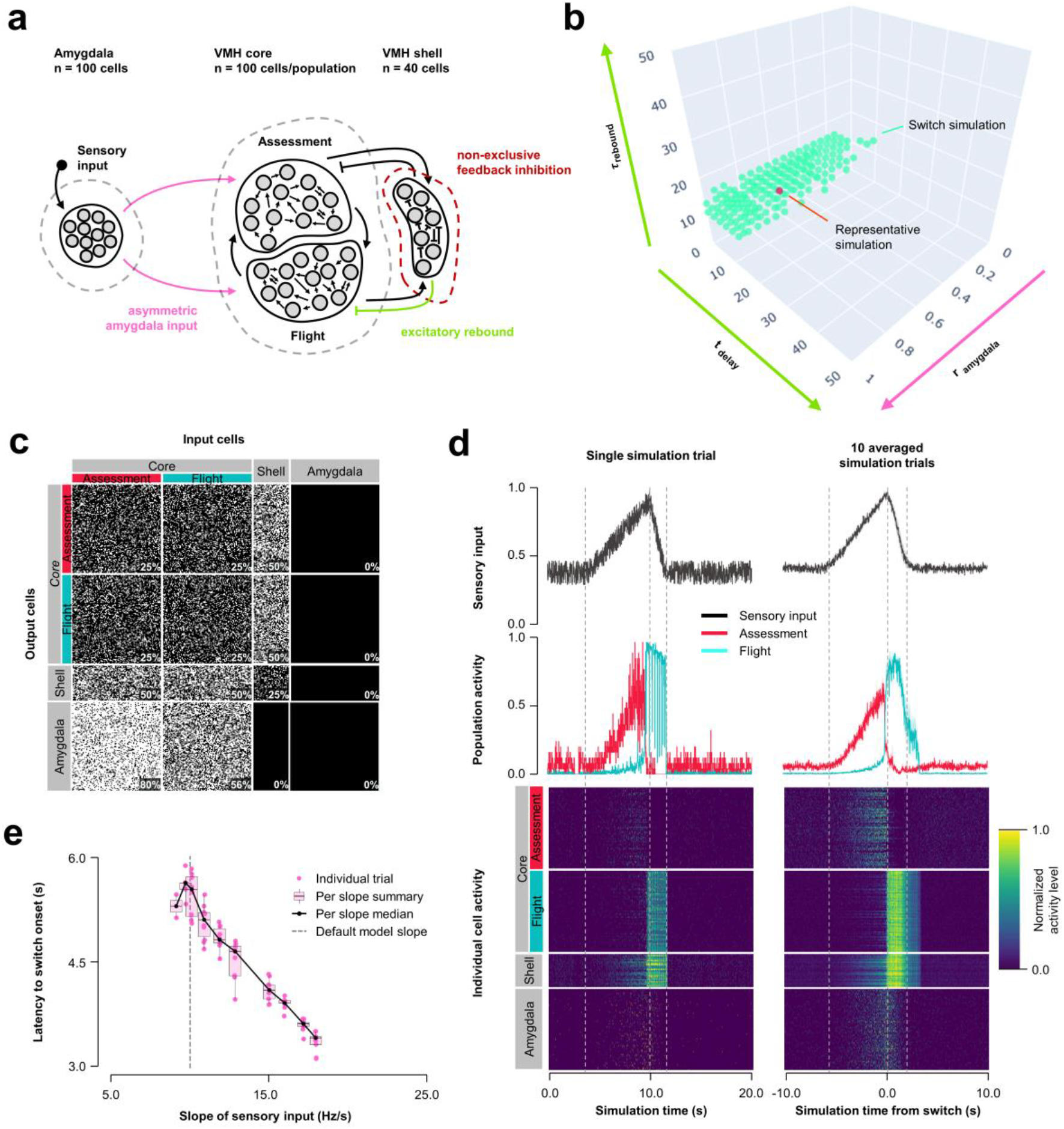
Model 4: Non-exclusive feedback inhibition. (**a**) Circuit structure with shell neurons joined into a single population (dark red) and stochastic core-shell and shell-core connectivity. As in Model 3, the delay time and membrane time constant for excitatory rebound in all inhibitory inputs onto Flight neurons (green) and the ratio (*r*_*input*_) of the probability of Amygdala-Flight/Amygdala-Assessment connectivity (pink) were systematically varied. (**b**) Systematic model exploration identified parameters that satisfy criteria for Assessment/Flight switching (turquoise, see **Methods**), where only simulation parameters that satisfy these criteria are shown. A selected representative simulation is indicated (orange). (**c**) Network connectivity matrix of the representative simulation showing connectivity probabilities between all neuron populations. (**d**) Network firing activity for the representative simulation from a single-trial simulation run (left) or a 10-trial averaged simulation run (right). Top row shows sensory input, middle row shows firing patterns of Assessment (red) and Flight (blue) populations, and bottom row shows firing activity of all individual cells, separated by population. All trials are centered around the time the switch occurs before being averaged. The vertical dashed lines mark the beginning and end of the assessment and flight phases of the sensory input (see **Methods**). (**e**) Plot of sensory input slope vs. latency of switching showing an inverse relationship between latency and sensory threshold. Note: only slope values for which a switch was observed are indicated.

## Discussion

Our computational circuit model of the mouse VMH was constrained by physiological properties derived from *in vitro* slice recordings of VMH neurons [23,26] and anatomical connectivity derived from *in vivo* tracing studies. [26,36,40] Our aim was to determine whether a relatively simple circuit architecture consisting of excitatory core neurons receiving excitatory sensory input and having excitatory and inhibitory feedback connections (**Fig 1–4a**) was sufficient to give rise to neuronal populations exhibiting sensory response patterns similar to the *Assessment*+ and *Flight*+ neurons identified by *in vivo* single unit recordings in behaving mice in response to a live predator [16] or social aggressor. [10] We found evidence that such a circuit can support the emergence of distinct neuron populations with reciprocal sensory input responses, and that these populations can exhibit abrupt switching activity that is thresholded to sensory input when excitatory slow-acting rebound properties are applied to feedback inhibition onto a sub-population of core neurons (Model 3-4; **Fig 3, 4**).

Several conclusions can be drawn from our findings. First, a simple circuit architecture of interconnected excitatory neurons receiving feedback inhibition was sufficient for the emergence of distinct populations of neurons with stable sensory ON and OFF response properties. A previous computational model of VMH circuitry exhibited similar sensory ON/OFF neuron populations [20] and in the mammalian striatum sparse lateral feedback inhibition has been shown to play a role in stabilizing ON and OFF firing states [41–45] suggesting that feedback inhibition may be a general circuit mechanism to establish stable reciprocal sensory response properties. However, unlike in striatum where the behavioral correlates of sensory ON/OFF neurons are not know, in VMH the switch between *Assessment+* and *Flight+* neuron firing is tightly associated with an approach-to-avoidance sensory-motor transformation [16] and thus establishes a testable correlation between behavior and circuit state transitions. Second, our models showed that a simple feedback inhibition architecture can give rise to switching activity that is thresholded to a gradually increasing sensory input. Such thresholding emerged either under specific unbalanced sensory input and feedback inhibition conditions (Model 1, **Fig 1**) or under balanced connectivity conditions that included unbalanced neuromodulatory rebound excitation (Model 3-4, **Fig 3-4**). Thresholded switching offers a mechanism by which accumulating sensory information about threat intensity could trigger the Assessment-to-Flight firing state transition in VMH and potentially engage downstream outputs to initiate escape behavior. It is noteworthy that reciprocally switching *Assessment+* and *Flight+* neurons are also found in the major downstream target of VMH, the periaqueductal gray (PAG), [16,46] suggesting that ancient instinctive defense pathways in the brain may have evolved as a hierarchical cascade of circuits that each allow sensory-to-motor transition decisions to emerge along the path from sensory input to locomotor output. A clue to the existence of such distributed sensory-motor decision control arises from the observation that feedback projections from PAG to VMH exert opposing excitatory and inhibitory effects, respectively, on *Assessment+* and *Flight+* cells in VMH [16] possibly modulating upstream decision nodes along the sensory-motor pathway. Third, the emergence of Assessment and Flight populations in our models persisted under conditions of equally weighted input and/or feedback connectivity (Model 2-4, **Fig 2-4**). Such a stochastic wiring of VMH core and shell neurons is consistent with developmental axon pathfinding processes guided by general target attraction and repulsion rules rather than those required for labelled-line connectivity and may provide benefits in the face of evolutionary pressure. [47]

Our models make several assumptions that may have biased our results. First, we did not include feedforward inhibition despite anatomical evidence to support its existence in projections from amygdala to VMH. [26,48,49] At present it is not clear what might be the impact of such feedforward inhibition on sensory responding of VMH core neurons, but we can conclude that feedforward inhibition is not necessary for the emergence of distinct Assessment and Flight neurons. Feedforward inhibition by VMH afferents is further complicated by the presence of parallel indirect amygdala input pathways to medial hypothalamus via pallidal structures, in particular bed nucleus of stria terminalis (BNST). [49–51] It has been proposed that such direct and indirect pathways allow for rapidly modulated go-no-go control of behavior as has been described in the basal ganglia. [50] Future models will need to explore the circuit properties afforded by such feedforward control in the medial hypothalamus. Second, we restricted our model to local VMH circuitry and did not attempt to include other nuclei that together form the highly interconnected medial hypothalamic defensive and reproductive systems and their immediate inputs and outputs. [4] For example, it has been shown that VMH projections to anterior hypothalamic nucleus are required for the production of defensive flight, [52] while the dorsal premammillary nucleus has been shown to support flight from both predators and conspecific threats. [17,53,54] Moreover, VMH is known to project to nuclei from which it receives projections [40] opening the possibility that multi-synaptic pathways among medial hypothalamic nuclei and their input and output structures could provide important additional circuit mechanisms for supporting the switching of *Assessment*+ and *Flight*+ cells in VMH.

We interpret the slow-acting excitatory rebound effect observed *in vivo* after sustained inhibition of VMH *Attack*-, but not *Attack*+ neurons, [23] as evidence that *Flight*+ and *Assessment*+ neurons may have distinct biophysical properties. In Model 3 we found that conferring excitatory neuromodulatory rebound to inhibitory inputs selectively onto Flight neurons was sufficient for the emergence of distinct Flight and Assessment neuron sensory responses and for their thresholded switching (**Fig 3**). Because Flight and Assessment neurons were otherwise uniform in this model, our findings demonstrated that slow-acting rebound excitation alone can confer the unique sensory response pattern of Flight neurons. Furthermore, they suggest that the selective expression of an excitatory neuromodulatory receptor on Flight neurons and the release of the cognate neuromodulator from inhibitory shell neurons could be a physiological mechanism sufficient to confer switching on VMH core neurons *in vivo*. VMH expresses several neuropeptides with excitatory metabotropic receptor signaling [20,28]; however, to date none of these has been implicated in defensive behavior. Notably, there is precedent in hypothalamic structures for the co-expression of GABA with neuropeptides that are released only under high frequency firing conditions. [55,56] Activation of such dual-neurotransmitter neurons at low frequency would be inhibitory, but at high frequency could be excitatory and provide the excitatory rebound to inhibitory synapses explored in our models.

Several of our findings present hypotheses that can be experimentally tested with existing *in vivo* circuit recording and manipulation tools. Simultaneous *in vivo* single unit electrophysiological recording and optogenetic stimulation (optrodes) could be used to test whether VMH *Assessment+* and *Flight+* neurons receive balanced amygdala inputs. A similar approach could be used to test whether VMH *Assessment+* and *Flight+* neurons receive inhibitory inputs from VMH shell with or without excitatory neuromodulatory rebound. Furthermore, our models make specific predictions about the sensory response properties of VMH shell inhibitory neurons that could be tested by cell-type specific recording. In particular, while our initial models (Model 1-3, **Fig 1–3a**) predict the presence of *Assessment+* and *Flight+*-like VMH shell neurons, our non-exclusive feedback inhibition model (Model 4, **Fig 4a**) predicts the existence of VMH shell neurons with a variety of sensory response properties, including hybrid *Assessment+* and *Flight+* properties (i.e. increasing during both approach and flight). Finally, our exploration of the connectivity properties that most effectively modulate the sensory threshold for Assessment/Flight switching points to a key role for feedback inhibition, rather than excitation in setting the approach-to-flight decision point. Thus, modulatory inputs onto VMH shell such as those that could be achieved by cell-type specific pharmacogenetic inhibition and activation should be highly effective at setting flight threshold and should have a predictable effect on sensitivity to threat. We expect that such experimental testing will in turn uncover new features to be incorporated in future computational models as part of a virtuous computational-experiment cycle.

## Supporting information

Table S1: All simulation parameters used.

## Author contributions

All simulated models and data analysis were carried out by R.R.; C.T.G. conceived the project and together with H.A. supervised the project; R.R., H.A., and C.T.G. designed the models and wrote the manuscript.

## Acknowledgments

We thank Maria Esteban Masferrer for providing access to the *in vivo* VMH recordings and for feedback on the model structures. The work was supported by EMBL.

