## Supplementary material for "Sensory-thresholded switch of neural firing states in a computational model of the ventromedial hypothalamus": Figures S1-4.

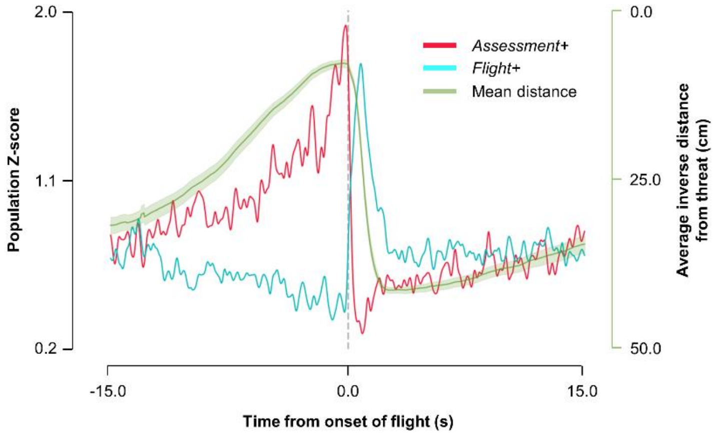


***Figure S1*. Firing state switch in VMH.** *In vivo* electrophysiology revealed an abrupt and reciprocal switch in firing states in the mouse VMHdm, as seen in the trial-averaged firing of representative *Assessment*+ (red, average N = 24) and *Flight*+ (blue, average N = 19) neurons (data reproduced from [16]). An additional trace and its corresponding y-axis (in green) show the mean and standard error of the inverse distance of the mouse to the threat during corresponding approach-avoidance behavior.

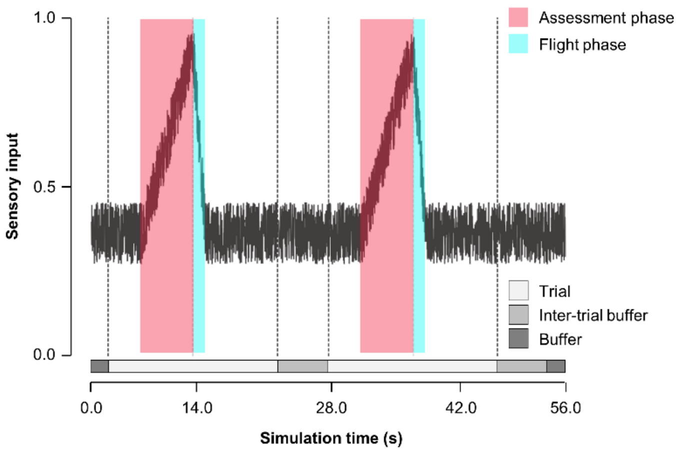


***Figure S2.*** **Sensory input function.** Noisy sensory input function $\lambda(t)$ used in our models to approximate the fluctuating sensory input mice receive during repeated approach toward and flight away from a stationary threat. Each approach-to-flight trial (two trials are shown here) consists of an approach (red, increasing sensory input) and subsequent flight (blue, decreasing sensory input) phase and is followed by an inter-trial buffer (yellow, stable sensory input) period, with the entire simulation starting and terminating with buffer periods (grey, stable sensory input).

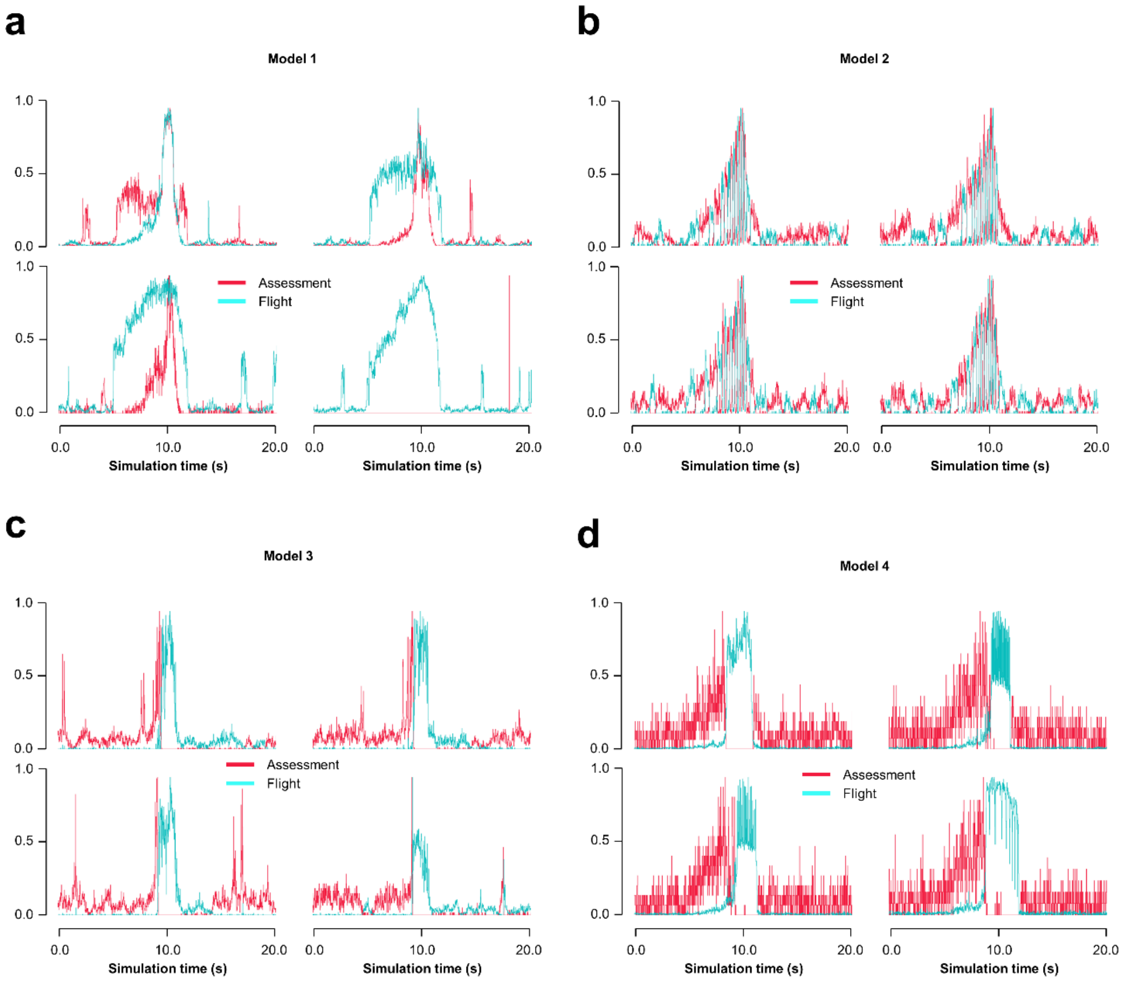


***Figure S3*. Model results with balanced sensory input.** Four example single-trial runs from the parameter exploration of each model with balanced sensory input ($r_{input} = 1$) and varying feedback inhibition, plasticity, and excitatory rebound parameters. Under these conditions Model 1 simulations (**a**) lose their ability to show a switch, while Model 2 simulations (**b**) still show intermittent burst firing and Model 3 (**c**) and Model 4 (**d**) simulations still show the switch.

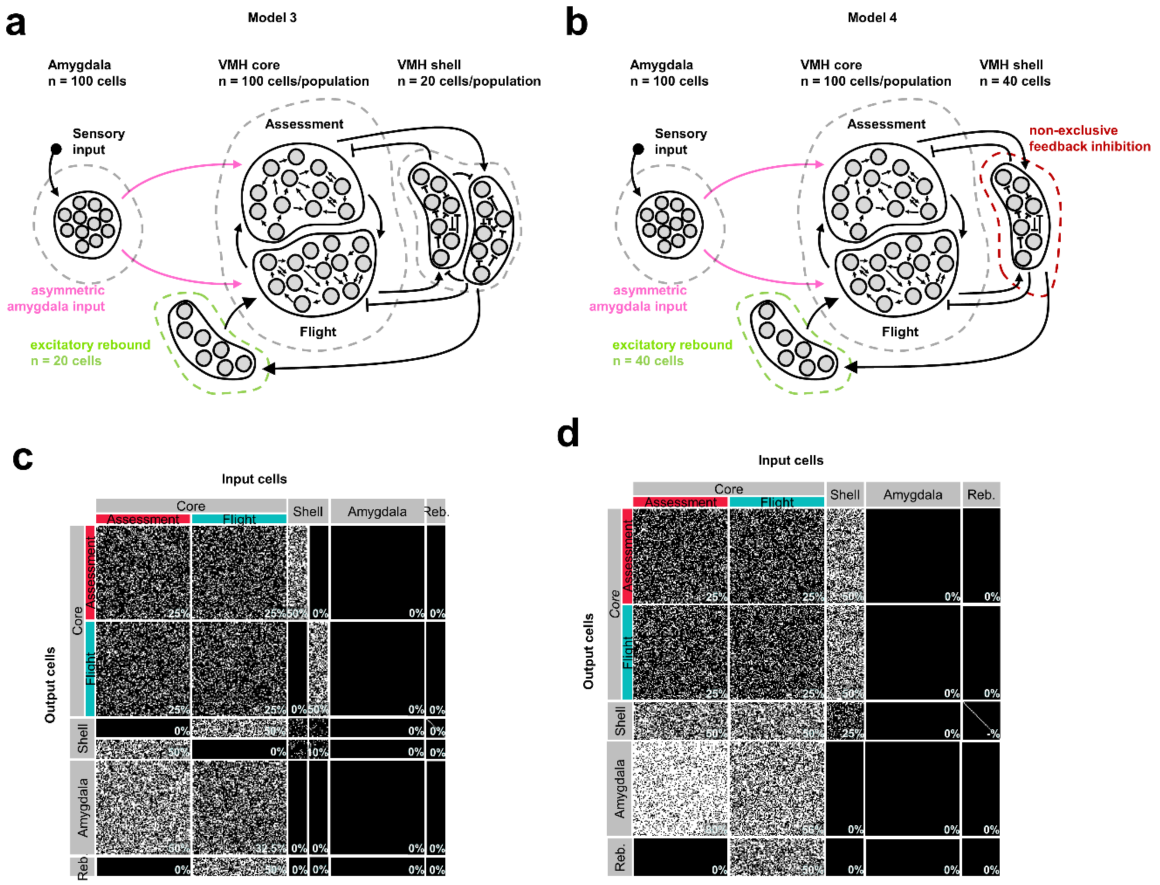


***Figure S4*. Implementation of excitatory rebound.** Circuit structure and connectivity matrix of (**a** and **c**, respectively) Model 3 and (**b** and **d**, respectively) Model 4 showing the added intermediate rebound population required to model inhibitory inputs with slow-acting excitatory rebound.
